# Enable, Empower, Succeed: Harnessing Open Science for Antimicrobial Resistance Containment

**DOI:** 10.1101/2024.02.20.580892

**Authors:** Luria Leslie Founou, Opeyemi U. Lawal, Armando Djiyou, Erkison Ewomazino Odih, Daniel Gyamfi Amoako, Stephane Fadanka, Mabel Kamweli Aworh, Sindiswa Lukhele, Dusanka Nikolic, Alice Matimba, Raspail Carrel Founou

**Affiliations:** Reproductive, Maternal, Newborn and Child Health (ReMARCH) Research Unit, Research Institute of the Centre of Expertise and Biological Diagnostic of Cameroon(CEDBCAM-RI); Bioinformatics & Applied Machine Learning Research Unit, EDEN Biosciences Research Institute(EBRI), EDEN Foundation, Yaoundé Cameroon; Antimicrobial Research Unit, School of Health Sciences, College of Health Sciences, University of KwaZulu-Natal, Durban 4000, South Africa; Canadian Research Institute for Food Safety, Department of Food Science, University of Guelph, Ontario, N1G 2W1, Canada; Virology, Mycology and Parasitology Laboratory, Postgraduate Training Unit for Health Sciences, Postgraduate school for pure and applied sciences, The University of Douala, PO Box 2701, Douala, Cameroon; Global Health Research Unit for the Genomic Surveillance of Antimicrobial Resistance, Department of Pharmaceutical Microbiology, Faculty of Pharmacy, University of Ibadan, Ibadan, Oyo State, Nigeria; Department of Pathobiology, University of Guelph, Guelph, Ontario, N1G 2W1, Canada; Mboalab Biotech, Yaoundé, Cameroon; Department of Population Health and Pathobiology, College of Veterinary Medicine, North Carolina State University, Raleigh, NC, United States; Computational and Integrative Biomedical Division, Faculty of Health Sciences, University of Cape Town, Cape Town, South Africa; Wellcome Connecting Science, Wellcome Genome Campus, Hinxton, Cambridge, United Kingdom; Antibiotic Resistance Infectious Diseases (ARID) Research Unit, Research Institute of Centre of Expertise and Biological Diagnostic of Cameroon (CEDBCAM-RI), Yaoundé, Cameroon; Department of Microbiology, Hematology and Immunology, Faculty of Medicine and Pharmaceutical Sciences, University of Dschang, Dschang, Cameroon

**Keywords:** Antimicrobial resistance, bioinformatics, capacity building, TtT, skills development, resource-constrained settings, Cameroon, Africa

## Abstract

Antimicrobial resistance (AMR) poses a significant threat to global health, particularly in Western sub-Saharan Africa where 27.3 deaths per 100,000 lives are affected, and surveillance and control measures are often limited. Genomics research plays a crucial role in understanding the emergence, spread and containment measures of AMR. However, its implementation in such settings is particularly challenging due to limited human capacity. This manuscript outlines a three-day bioinformatics workshop in Cameroon, highlighting efforts to build human capacity for genomics research to support AMR surveillance using readily accessible and user-friendly web-based tools. The workshop introduced participants to basic next-generation sequencing concepts, data file formats used in bacterial genomics, data sharing procedures and considerations, as well as the use of web-based bioinformatics software to analyse genomic data, including *in silico* prediction of AMR, phylogenetics analyses, and a quick introduction to Linux© command line.

We provide a detailed description of the relevant training approaches used, including workshop structure, the selection and planning, and utilization of freely available web-based tools, and the evaluation methods employed. Our approach aimed to overcome limitations such as inadequate infrastructure, limited access to computational resources, and scarcity of expertise. By leveraging the power of freely available web-based tools, we demonstrated how participants can acquire fundamental bioinformatics skills, enhance their understanding of biological data analysis, and contribute to the field, even in an underprivileged environment.

Our findings highlight the effectiveness of this training approach in empowering local researchers and bridging the bioinformatics gap in genomics surveillance of AMR in resource-constrained settings. Building human capacity for genomics research globally, and especially in resource-constrained settings, is imperative for ensuring global health and sustainable containment of AMR.

**Data summary:** The authors confirm that all supporting data, code and protocols have been provided within the article or through supplementary data files.

**Impact Statement:** Antimicrobial resistance (AMR) is a major global health threat, especially in Western sub-Saharan Africa, where 27.3 deaths per 100,000 lives occur. Genomics research play an instrumental role for understanding AMR’s emergence, spread, and containment measures. However, its implementation in these settings is challenging due to limited human capacity. A three-day bioinformatics workshop in Cameroon aimed to build human capacity for genomics research using web-based tools. Participants were introduced to next-generation sequencing concepts, data file formats, data sharing procedures, and web-based bioinformatics software for analysing genomic data.

The workshop aimed to overcome limitations like inadequate infrastructure, computational resources, and expertise scarcity. The findings show the effectiveness of this training approach in empowering local researchers and bridging the bioinformatics gap in genomics surveillance of AMR in resource-constrained settings.

## Introduction

Antimicrobial resistance (AMR) is a quintessential One Health issue that has multidimensional implications for humans, animals, and the environment (1). AMR caused 1.27 million deaths and was estimated by predictive models to be associated with nearly 5 million deaths globally in 2019 (2). It is predicted to become the leading cause of mortality, claiming 10 million lives yearly with an estimated cost of USD$100 billion on a global scale by 2050, if nothing is done to sustainably contain it (1). With the advent of whole genome sequencing (WGS), the increasing affordability of sequencing technologies, and the expansion of genomic studies generating large datasets from various sources, bioinformatics has become instrumental in gaining new insights into the complex molecular mechanisms and transmission dynamics of AMR. Informatics has revolutionized the fields of biology and microbiology by providing powerful tools and techniques for analysing and interpreting complex biological data with granular resolution (3). More specifically, bioinformatics has been recognized as a crucial field that can help in the development and design of vaccines, medicines, and diagnostics tools, as well as in the implementation of tailored prevention measures to contain AMR (4-6). Nowadays, bioinformatics skills serve as building blocks for scaling up genomics surveillance of AMR (7, 8). Open science is increasingly acknowledged as a critical accelerator for bioinformatics and scientific knowledge accessible to everyone and in an inclusive, equitable and sustainable manner.

However, the adoption of bioinformatics approaches in resource-constrained countries, particularly in Africa, is hindered by several challenges, including limited access to computational infrastructure, a shortage of trained personnel, and financial constraints, among others (9). In Cameroon, the human workforce with the required bioinformatics skillset is scarce, with few universities offering such a curriculum. To acquire bioinformatics skills, many researchers or professionals rely on online courses and workshops or seek training outside the country or continent through migration. The scarcity of bioinformatics expertise and training has a cascading effect on the establishment of large-scale and high-throughput genomics projects that support the sustainable containment of AMR in the country. Consequently, there is a pressing need for innovative approaches to overcome these barriers and empower researchers in the country to harness the potential of bioinformatics. Although there are increasing capacity development initiatives spreading across the continent, these rely on input from few groups or institutions based in Africa or abroad. To address this, the Research Institute of the Centre of Expertise and Biological Diagnostic of Cameroon (<u>CEDBCAM-RI</u>) developed and carried out, to the best of our knowledge, the first bioinformatics workshop focusing on bacterial AMR in Cameroon.

The goal of this workshop was to initiate building of human capacity for the analyses of genomics data to support AMR surveillance in Cameroon. The vision inherent to this three-day workshop was to introduce researchers or professionals to the field of bioinformatics with focus on AMR and equip them with the minimum bioinformatics skillset required to support bacterial genomics research in Cameroon. Specifically, by using learner-centred training approaches, it aimed to (i) impart participants with knowledge of the underlying principles of next-generation sequencing (NGS) and bioinformatics analyses, (ii) equip them with bioinformatics skills to analyse and interpret bacterial genomics data using free web-based tools, and (iii) establish a network of scientists working towards AMR surveillance and containment. The workshop was entitled “Bioinformatics applications in genomics surveillance of bacterial antimicrobial resistance” formed part of the CEDBCAM Bioinformatics Empowerment (CBE) initiative. This article summarizes the development and implementation of this workshop and showed how online-based tools were leveraged to overcome resource limitations and strengthen the genomics research landscape in Cameroon. By sharing our experiences, we hope to provide a blueprint and inspiration to other resource-constrained settings facing similar challenges.

## Methods

A unique and forward-thinking approach that combines education, relevant training approaches, utilization of freely available and widely used bioinformatics web-based software, and collaborative efforts were used to empower local scientists. Figure 1 provides an overview of the vision inherent to this workshop. The workshop targeted postgraduate students, and professionals with background in biology or health sciences.

**Figure 1.**
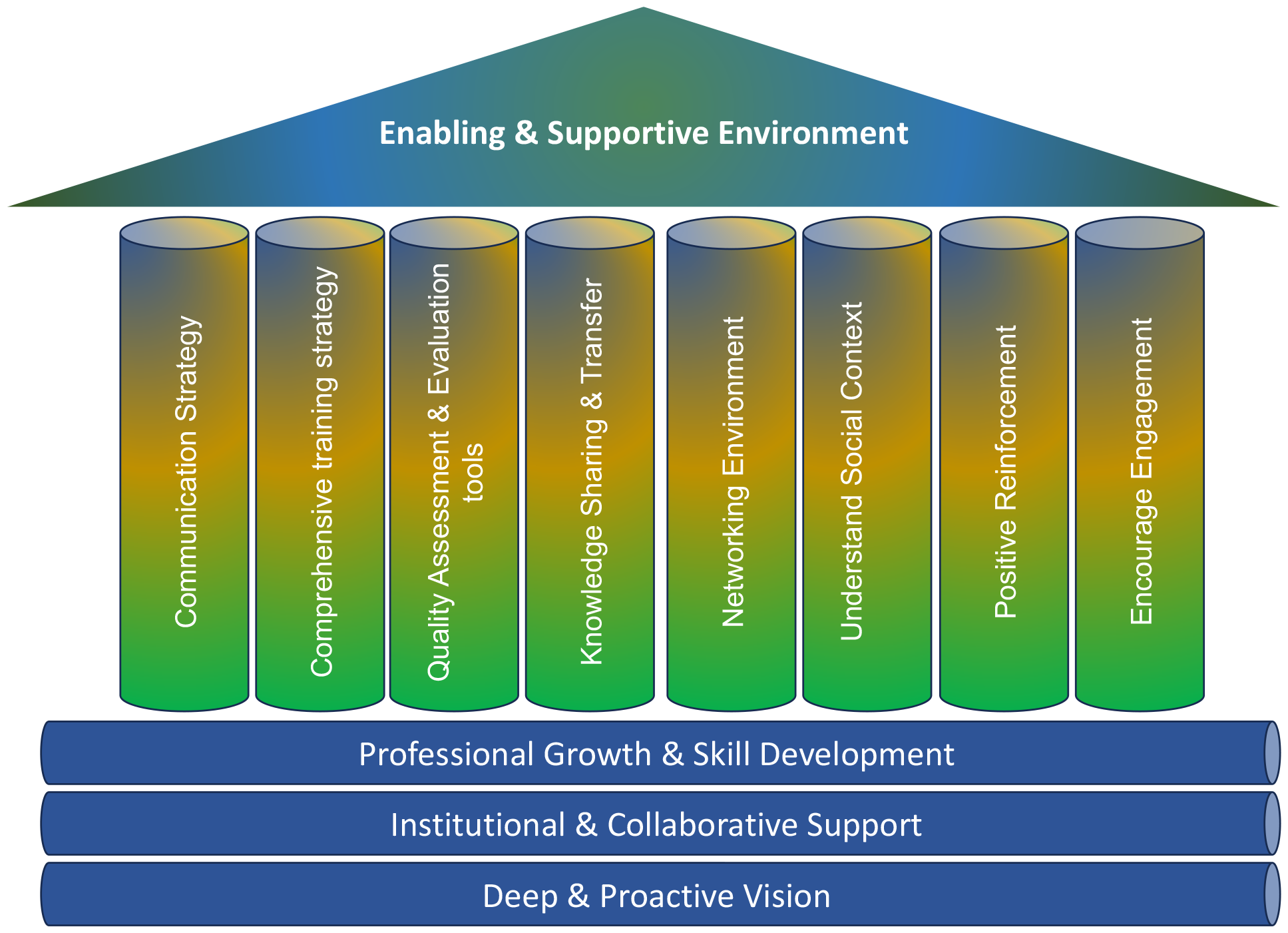
Overview of the workshop’s vision.

### Workshop Preparation

The workshop was designed by leading organizers and researchers of CEDBCAM-RI. The team consisted of LLF, SF, and RCF. As a group, we held a workshop to identify challenges and discuss solutions to implement the training programme within the constraints of inadequate local bioinformatics expertise, administrative, infrastructure and logistical limitations. The outcome of these discussions was used to tailor the training programme (Table 1). Administrative support was obtained from <u>CEDBCAM-RI</u> executive board to ensure the success of the workshop. The workshop duration and schedule were tailored to accommodate the participants’ and facilitators’ availability and ensure effective knowledge transfer. A minimal registration fee of USD $100 was requested from participants for catering purpose. This fee was fully or partially waived for selected participants with no or limited financial resources thanks to support received from some partners. CBE workshop was advertised through social media platforms.

**Table 1.** Summary of challenges identified, and solutions implemented during the preparation of the workshop.

| Priority Level | Description of challenges | Solutions and risk mitigation measures implemented |
| --- | --- | --- |
| **High** | Lack of trainers | We made effort to invite some experienced trainers from H3ABioNet, Wellcome Connecting Science, and SeqAfrica as well as collaborative institutions like Wellcome Sanger Institute. |
|  | High cost associated with on-site workshop | Basic internal funding was provided by CEDBCAM-RI<br>Requested small support from suppliers and collaborating institutions.<br>A minimum registration fees was required to attend the training |
|  | Timely development of the course materials | Preparatory meetings were planned and carried out.<br>Tasks were assigned and each facilitator was responsible of his/her modules/activities.<br>Communication was key with group created on social media application and with repetitive email. |
| **Medium** | Support from CEDBCAM-RI administrative board | The support from CEDBCAM-RI was requested during the end of year general assembly defining the key objectives and activities of the next year. Having forward thinking management with long-term plan and vision helped securing support for the workshop. |
|  | Language barrier – Most of the participants were French native speakers while half of the facilitators were English native speakers. | Guest speakers and facilitators were informed on the main native language of participants.<br>On-site facilitators were bilingual and serve as translators for the speakers and facilitators as well as for the participants as far as it was needed.<br>Course materials were provided before, and recording were made available to allow participants to watch again the lectures and discussion sessions. |
|  | Network issue | Although the CEDBCAM-RI has good and unlimited internet network, alternative modems were installed as backup.<br>Participants were also requested to come along with their personnel modem as from our experience some participants may have computers not taking any of the existing network.<br>Video recordings of the guest speakers and some lecturers were sent before the workshop and played during the workshop |
|  | Infrastructure – Poor energy supply in the country may disrupt the course. | A generator was available at CEDBCAM-RI and more uninterruptible power supply were added. |
| **Low** | Lower than expected participation rates | We emphasized communication and expanded the target from PhD students or holder to Master students, and professionals |
|  | Dropouts of some participants | A restricted number of participants (maximum of 15) was selected |
|  |  | and close monitoring and follow up for the trainee was conducted |
| **Very Low** | Lack of computer room on site which may not allow participants to fully enjoy the course if they lack a good computer. | A large conference room was used for the training within the CEDBCAM-RI.<br>The online-based tools require a normal computer and network. Participants were requested to come along with their personal laptop |

### Risk analysis and contingency plan

As this was the first time this type of training as being held, a risk analysis and contingency plan was developed based on an impact/effort matrix including four categories: major tasks, quick wins, thankless tasks, and fill-ins. The major tasks were those requiring considerable efforts from organizers, course developers and facilitators, and teaching assistants to achieve impactful training. The quick wins tasks were those having high impact on the workshop but requiring low effort or efforts from lead organizers only such as getting administrative support, addressing language barrier, and ensuring stable network and power supply. The thankless tasks were those requiring high effort from lead organizers and teaching assistants but having a lower impact on the workshop, including catering, advertisement, agenda, and infrastructure setting. Finally, fill-ins were unexpected tasks with low impact and low effort that were added during the running of the workshop. A mitigation/contingency plan was developed with high, medium, low, and very low priority measures (Figure 2). The workshop organizers collaborated with international experts from regional networks such as H3ABioNet network (https://www.h3abionet.org/) and SeqAfrica (https://antimicrobialresistance.dk/seqafrica.aspx) consortium.

**Figure 2.**
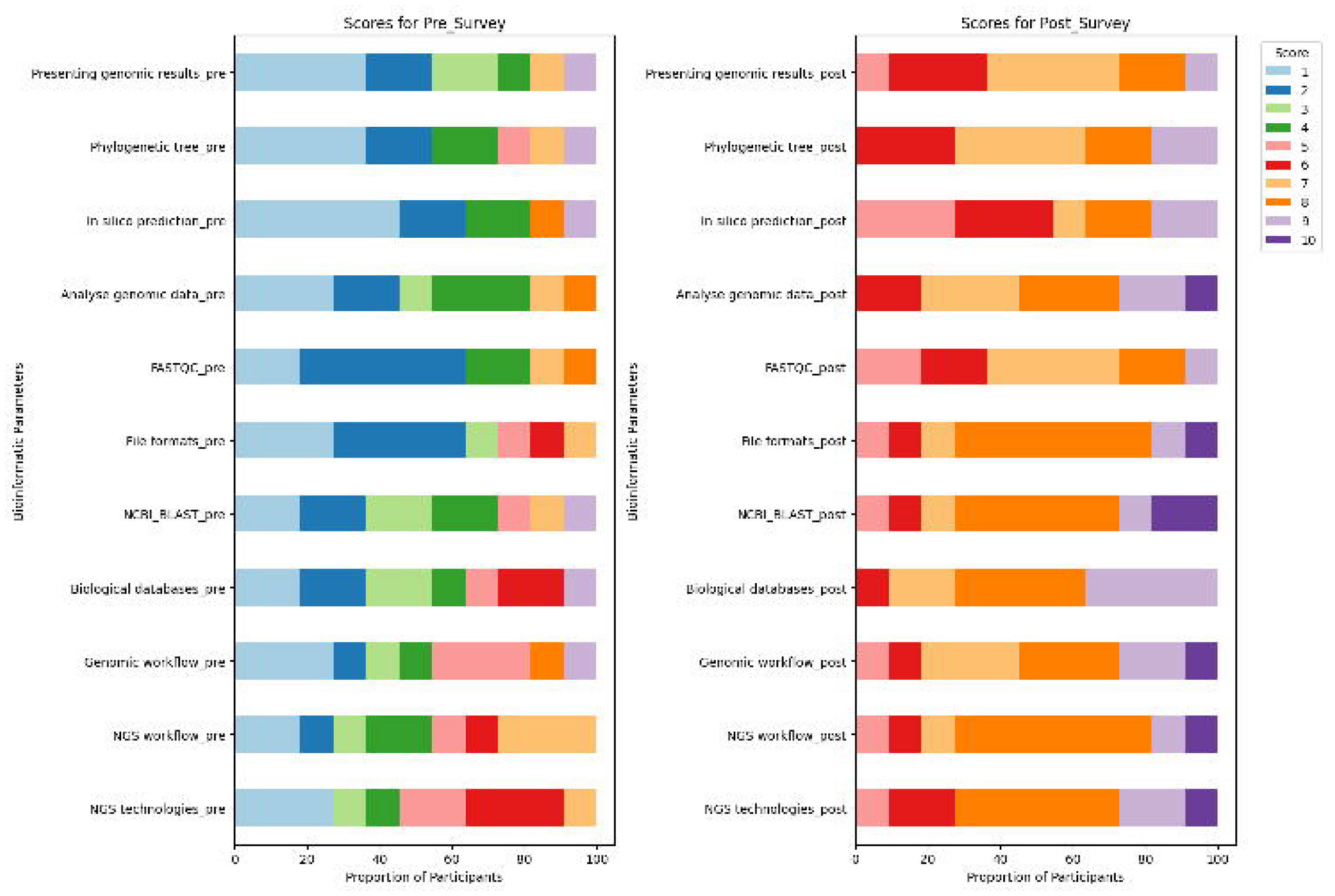
Impact/Effort matrix and mitigation/contingency plan.

### Participant Selection and Recruitment

A call for participation was launched one month before the workshop with questions targeting prospective participants’ current educational level, knowledge of the basics of genetics and genomics, as well as current and future research goals. A selection committee was constituted to assess all the applications based on predefined selection criteria. Participants were selected based on their interest in AMR, their background in biology-related field, understanding of the importance of genomics in the surveillance of AMR, and their potential to apply immediately the acquired knowledge in their research or respective institution. Recruitment efforts targeted researchers and individuals from academia, research, industry, or clinical settings, to ensure that the workshop reached those who would benefit the most from enhanced bioinformatics skills.

### Curriculum development

The workshop curriculum was created through collaboration among bioinformatics experts and educators, using an outcome-based approach to training design. The workshop design started with setting off the learning outcomes using an outcome-driven approach. With a target audience consisting of healthcare professionals and researchers, principles of adult training were incorporated into the design of sessions, and activities were based on an active learning approach that included hands on activities and diverse ways of learners’ engagement, such as group work, peer discussions, and interactive ways of formative assessment. These approaches were directly informed by a genomics training design framework developed by Wellcome Connecting Science (https://wcscourses.github.io/T3connectResources/) which assumes a training framework adaptable to various contexts and different domain specific knowledge training. A member of CEDBCAM-RI team was a trainee of the WCS/H3ABionet Train the trainer course (https://coursesandconferences.wellcomeconnectingscience.org/event/train-the-trainer-course-design-and-delivery-for-bioinformatics-trainers-virtual-20221122/) and leveraged the TtT facilitator expertise to draft the curriculum through several rounds of discussions and peer review, which helped with the integration of the training framework used into the specific context of the CEDBCAM workshop, called CBE 2023.

After mapping the needs and levels of the target audience, content and activities were developed consisting of bioinformatics concepts, tools, and interpretation. The content was structured to introduce participants to fundamental concepts, including NGS technologies, sequence analysis workflow, sequence alignment, *in silico* prediction, phylogeny and an introduction to Linux© and data sharing. Each separate module further followed more detailed learning outcomes. The activities included lectures and hands-on exercises using web-based tools such as the Linux emulator (https://www.cygwin.com/). Group discussions offered opportunities for peer experience and instructor-experts knowledge sharing and served as formative assessment for the participants. The duration of each activity (and each module) was assessed in advance and agreed to by facilitators. Where facilitators assisted remotely, video recordings of course material were provided as explained via the acceptance letter. To allow for assessment of the learning outcomes, adequate learners’ assignments were set. Those were of practical nature, corresponding to the introductory but applicative nature of the course (as per the Blooms pyramid of cognitive development levels, also used for describing of the LOs). The modules implemented are provided in Table 2.

**Table 2:** Modules, objectives, and learning outcomes of the workshop.

| Modules | Summary objectives | Learning outcomes |
| --- | --- | --- |
| **Introduction to genomics and bioinformatics** | - The role of genomics and bioinformatics<br>- Application of bioinformatics to infectious diseases and AMR containment<br>- To recognise the job opportunity related to this field | - Define key theoretical concept of bioinformatics<br>- Recognise NGS technologies and NGS data file format<br>- Understand the sequencing workflow<br>- Use biological databases and resources to search and retrieve file |
| **Basic Local Alignment Tool (BLAST)** | - Sequence alignment theory and applications<br>- BLAST theory and practice<br>- Overview of primer design | - Interpret the information conveyed in NCBI BLAST search outputs and infer their significance<br>- Examine the annotations of reported matches and their provenance<br>- Export BLAST results |
| Online-based tools for bacterial genotyping | - Analyse the quality of genomic data with fastqc<br>- Explore web-based tools as a method to access publicly available genomes and analyse genomic data | - Perform quality check with fastqc<br>- Browse publicly available genomes and associated metadata.<br>- Use online tools to analyse genome sequence data<br>- Interpret output of *in silico* prediction |
| **Introduction to phylogenetics and phylogenetic analysis** | - Principles of phylogenetics<br>- Phylogenetic tree construction, interpretation, and visualisation | - Understand the key principles of phylogeny<br>- Use Patric for retrieval of available genome<br>- Construct a tree with snp-tree of RGI<br>- Visualise a phylogenetic tree with MicroReact<br>- Interpret a phylogenetic tree |
| **Introduction to Linux** | - Introduction to Unix, file structure and navigation<br>- Basic Linux commands<br>- Navigation in Linux directories<br>- Basic manipulating file commands | - Understand the Linux file structure<br>- Understand the command line structure and learn basic commands<br>- Create, access files and directories and navigate through them<br>- Read files content and extract information from them<br>- Run a pre-written Linux script |

### Selection of Web-Based Tools

Careful consideration was given to the selection of web-based tools to ensure their suitability for the level of participants and accessibility. Criteria included free availability, low computational and connectivity requirements (i.e. low internet bandwidth require to analyse data), user-friendly interfaces, relevance, and suitability in the field of AMR surveillance and availability of necessary bioinformatics functionalities. We sought to bring participants from the most rudimentary web-based tools, Basic Local Alignment Search Tool (BLAST; (https://blast.ncbi.nlm.nih.gov/Blast.cgi) to some of the most advanced online tools, CSIPhylogeny (https://cge.food.dtu.dk/services/CSIPhylogeny/), PathogenWatch (https://pathogen.watch/) and Bacterial and Viral Bioinformatics Resource Center (https://www.bv-brc.org/). The full list of online tools providing comprehensive functionality while minimizing the need for local infrastructure is provided in Supplementary Table 1.

### Running of the Workshop

The CBE workshop was conducted over three days from 22-24 February 2023 from 8.00 am to 6.00 pm (GMT+1). Physical attendance and completion of assignments were mandatory. The workshop was fully supported by CEDBCAM-RI internal funds. The agenda of the workshop is presented in Supplementary Table 2.

### Evaluation

To assess the effectiveness of the workshop, both formative and summative evaluations were employed. Formative evaluation involved continuous feedback collection throughout the workshop to gauge participants’ understanding, engagement, and satisfaction. Furthermore, group projects summarizing all aspects taught were assessed at the end of the workshop, to evaluate the delivery of learning outcomes. Learners reported their confidence in gaining skills and using the tools taught in pre- and post-workshop surveys, thus providing an indication of how much was learned during the workshop. Participant feedback surveys were also administered to gather qualitative data on the workshop’s impact and areas for improvement.

## Results

### Understanding barriers to implementation

During the preparation phase, several challenges were identified, and solutions proposed as outlined in Table 1. The most important challenge was the scarcity of bioinformatics experts specialised in AMR surveillance in the country. Another challenge was the financial constraints associated with any on-site training activities. Language barrier was another difficulty identified given that most participants speak French. Poor energy supply and unstable network throughout the country were structural and infrastructural challenges identified.

### Participants’ socio-demographic characteristics

From the 86 individuals who applied for the workshop, 11 participants from various institutions and origins were selected. Most participants were from University of Yaoundé I (58%) and PhD students (83%). The participants came from four out of the ten regions of Cameroon, and one participant was from Chad. Only four out of the eleven participants were female (36% vs 64%). The median age of participants was 29 years (IQR: 8.5; range: 23 – 39).

### Skill development and knowledge retention

Before and after the workshop, each participant completed a confidence survey as part of the summative evaluation. In summary, prior to the workshop, most participants (7/11; 64%) acknowledged that they lacked confidence in their ability to identify next-generation sequencing (NGS) technologies or NGS workflows, with a score of ≤5. Nine out of eleven participants (81%) expressed lack of confidence in their ability to identify and use NGS data formats and NCBI Blast for analysis. Furthermore, most participants (9/11; 81%) expressed a lack of confidence in their ability to analyse genetic data using web-based bioinformatics tools, with participants having a score ≤4 and 33% of them had a score of 1. Regarding the interpretation of *in silico* predictions of antibiotic resistance genes (ARGs), a similar pattern was noted, with 81 percent (9/11) of participants scoring ≤4, of whom 45 percent (5/11) scored 1. Nine out of eleven participants (81%) expressed lack of confidence in their ability to communicate genetic data at a conference (Figure 3).

**Figure 3.**
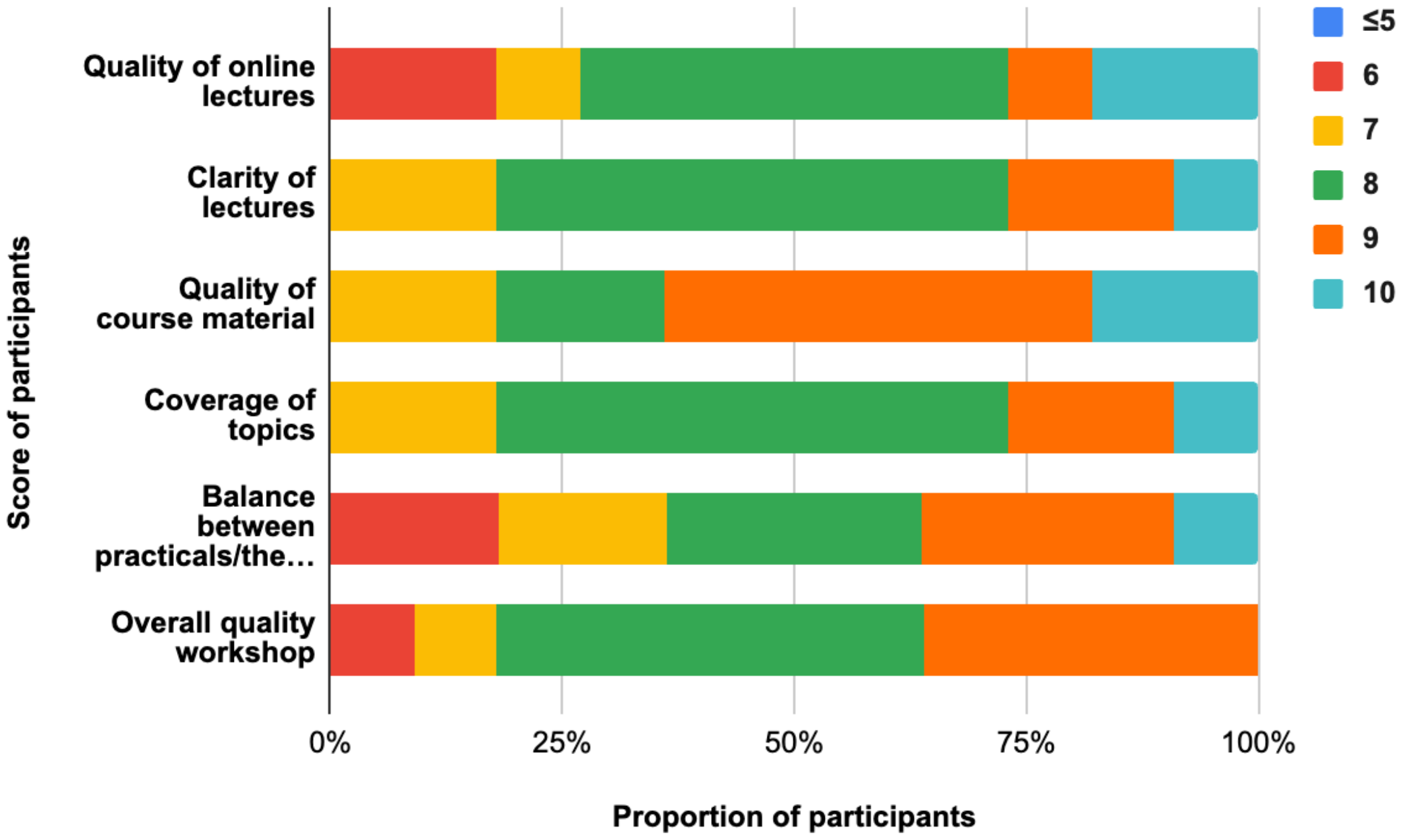
Summary of pre- and post-workshop survey results.

In contrast, after the workshop, a substantial increase in participants’ confidence in bioinformatics knowledge and skills was observed. Briefly, all participants were extremely confident in recognising NGS technologies or the NGS workflow with a score ≥5 for all participants out of which 55% (6/11) and 36% (4/11) had a score of 8 for NGS technologies and NGS workflow, respectively. In addition, all participants were confident in using NCBI BLAST for data analysis with a score ≥5 among which 45%, 9% and 18% had a score of 8, 9, and 10, respectively. A similar observation was made for the use of web-based bioinformatics tools for data analysis and presentation of genomic results at a conference. The great majority of participants reported increased confidence in interpreting *in silico* prediction analysis, with 81% (9/11) having a score >5 (Figure 3).

### Participant feedback and engagement

Participant feedback indicated an important level of satisfaction with the workshop content, delivery, and overall experience. The interactive nature of the workshop, combined with the practical exercises using freely available web-based tools, significantly enhanced participants’ engagement and understanding. Participants appreciated the helpfulness of the online lectures of the workshop with all of them giving a score ≥6 of which 45% (5/11) gave a score of 8 (Figure 4). Likewise, all participants valued the course materials provided, with an overall score ≥7, and 54% (6/11) of them reporting a rating of 9. The CBE workshop was concluded by a ceremony of hand-over certificate during which all participants affirmed that they would like to undertake the intermediate level of the workshop. They also expressed enthusiasm for incorporating bioinformatics approaches into their current and future research projects and appreciated the opportunity to network with peers facing similar challenges.

**Figure 4.**
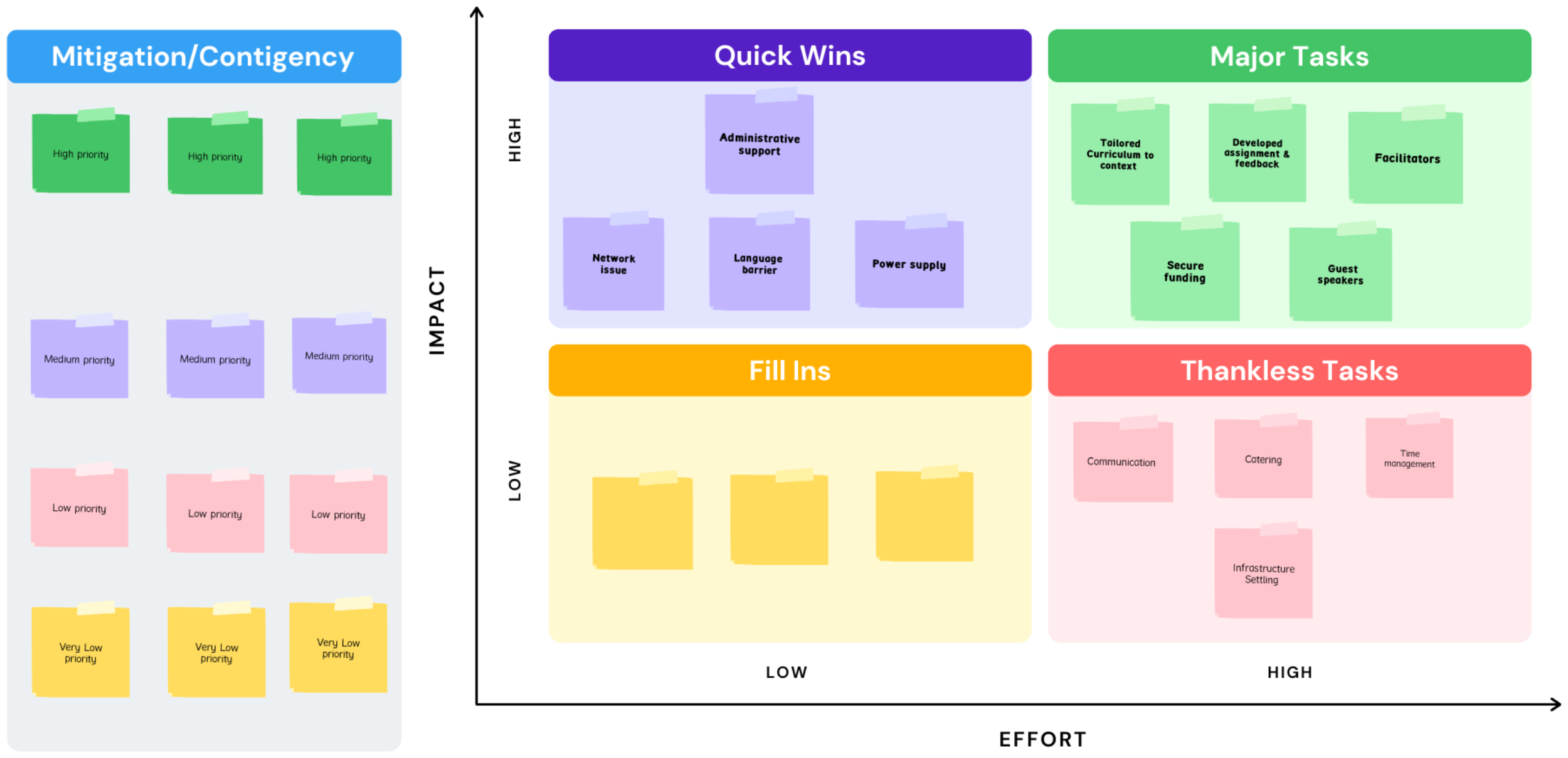
Evaluation of workshop.

## Discussion

Genomics offers invaluable insights into the understanding and implementation of containment strategies for AMR, a global public health crisis that compromises the efficacy of existing antimicrobial therapies. In resource-constrained settings, notably those in Africa, such as Cameroon, the challenges in combating AMR are exacerbated due to limited surveillance capabilities, inadequate laboratory infrastructure, and a shortage of well-trained personnel. The pre- and post-workshop surveys demonstrated significant progress in equipping participants with essential bioinformatics skills and knowledge using web-based tools as depicted in a significant enhancement in their confidence levels across various aspects of genomics and bioinformatics.

Studies conducted in similar resource-constrained settings consistently identified the shortage of bioinformatics expertise as a significant barrier to genomic surveillance efforts (10-12). The case of our CBE workshop in addressing this challenge resonates with efforts of various organizations and initiatives, such as H3ABioNet, Eastern Africa Network of Bioinformatics Training (EANBitT; http://cbid.icipe.org/apps/eanbit/), Collaborative African Genomics Network [CAfGEN (9) and SeqAfrica, recognizing the imperative need to strengthen bioinformatics skills in Africa (13, 14).

Compared to traditional in-person workshops, our hybrid-based training approach, where trainers had the opportunity to provide aid in-person or remotely exhibited distinct advantages, including facilitator location flexibility, and reduced financial burdens associated with travel and accommodation. It also provided continuous access to workshop materials and resources. Additionally, the utilization of freely available web-based tools proved to be a viable solution for sowing seeds towards AMR genomic surveillance in resource-constrained settings. These tools offered comprehensive bioinformatics functionality without requiring local computational resources, thus enabling participants to acquire essential skills regardless of institutional limitations.

Application of the TtT framework proved viable in responding to training needs in our specific context and audience and was well adaptable for training in the AMR specific knowledge and skills domain. Ensuring the sustainability of capacity building efforts is crucial for long-term success. To ensure the sustainability of the workshop’s impact, strategies were implemented to foster a supportive community of practice among participants. Group discussions, interaction with senior peers, mentorship, and collaboration opportunities were established to facilitate ongoing learning and knowledge sharing. Sensitising participants to pursue in their learning path and becoming trainers themselves further enhanced the long-term impact of the workshop, as they could disseminate their knowledge within their local contexts. The importance of regional and international collaborations in supporting ongoing research and surveillance activities was also emphasized. Sustaining human capacity, including continuing training, mentorship programs, and integration of genomics research into local academic and healthcare systems is imperative for sustainable containment of AMR (9).

Acknowledging the study’s limitations, the need for further expansion and improvement of similar initiatives is evident. First, while our workshop highlighted the potential of online tools, we acknowledge that relying on internet connectivity can be a limiting factor in resource-constrained settings. A potential avenue for future exploration is a conventional fully on-site workshop with computers coupled with software working offline. This approach would mitigate connectivity challenges and enhance the efficiency of bioinformatics analyses. Second, the short duration of the workshop, though effective in introducing fundamental concepts, leaves room for more extensive training. Long-term programs could delve deeper into advanced topics, fostering a greater depth of expertise among participants. Such extended training would align with the needs of researchers aiming to undertake large-scale genomics projects to contain antimicrobial resistance. Third, despite the commitment of our online trainers and senior experts to transfer their knowledge remotely, physical presence of both trainers and peers is acknowledged to have a higher impact than remote interactions. This is especially true given the context of the participants who raised valid concerns during discussions with senior peers “*how should we do genomics or vaccine prediction when in our hospitals where we are not able to do conventional bacteriology culture*;” “*how to think about sequencer when we do not have stable electricity*.” These statements revealed the mindset of some of our participants who despite agreeing with their peers, could not see how to reach their goals in their current environment. Physical contact and face-to-face discussion will have provided room to senior peers for high-level motivation, reinforcing inspiration, and self-belief messages to strengthen the participants’ ability to change the status quo. Fourth, language barrier is a crucial element to enable access to quality bioinformatics education. Whilst numerous efforts were made to facilitate understanding of participants, including sharing of resources and live translation from French to English and vice-versa, it is noteworthy to mention that efforts needed to understand the speakers may have impeded participants’ concentration.

As such, building on our successes and addressing the limitations we faced, we propose some recommendations for future capacity building initiatives:

1. **Leveraging local cluster infrastructure:** Web-based tools are adequate for acquiring fundamentals bioinformatics skillset, but for large-scale and more complex bioinformatics analysis, cluster infrastructures are crucially needed. In resource-constrained settings, partnering with local institutions to establish and maintain cluster infrastructure for bioinformatics analyses can provide more reliable and sustainable access to computational resources, reducing reliance on fluctuating internet connectivity. More stable internet connectivity is required to both run (ensure uptime) and use (connect and transfer data) clusters.
2. **Long-Term Training:** Explore the implementation of longer-term bioinformatics training programs to offer participants a more comprehensive skill set that encompasses advanced topics and would allow participants to become proficient in handling complex genomic data. The CafGen initiative in Uganda and Botswana is a good example (9).
3. **Evaluation and Impact Assessment:** Conduct rigorous evaluations of workshop outcomes, including long-term impacts on participants’ research and careers. This evaluation will provide valuable insights into the effectiveness of such initiatives and guide improvements.
4. **Use TtT framework:** integrate efficient training approaches with domain specific training to enable capacity building by forward teaching and training.
5. **Promote inclusivity:** Considering the main language of the audience is of paramount importance, especially in Africa where multiple languages coexist. Inclusivity suggests considering training initiatives that account for the languages spoken by the audience or minimize the gaps between participants. Acknowledging the linguistic diversity in Africa, with 21 of the 54 countries having French as an official language, is crucial.

## Conclusion

Building a human workforce for genomic surveillance of AMR can be initiated with as simple as a bioinformatics workshop using freely available web-based tools. By recognizing our local challenges and leveraging online bioinformatics tools, we have demonstrated the feasibility of developing human capacity for genomics research to support AMR surveillance in resource-constrained settings. Our bioinformatics workshop in Cameroon can serve as a promising model for capacity building in challenging environments. To ensure sustainability of AMR containment efforts, it is imperative to replicate and expand such training initiatives on a larger scale and for an extended duration.

## Acknowledgments

We would like to express our gratitude to the keynote speakers Prof. Stephen D. Bentley and Prof. Sabiha Yusuf Essack for their time and support in implementing this workshop. We also thank all the participants who actively engaged in the workshop and provided valuable feedback. We extend our appreciation to the instructors, organizers, and institutions that supported the development and implementation of this workshop.

## Conflict of interest

The authors declare that there are no conflicts of interest.

## Author contributions

**LLF**: conceptualization, funding acquisition, investigation, methodology, formal analysis, data curation, project administration, writing – original draft. **OUL, AD, EEO, SF, MKA**: investigation, data curation, methodology, writing – review and editing. **DGA**: investigation, data curation, methodology, formal analysis, writing – review and editing. **SL, DN, AM:** project administration, resources, supervision, writing – review and editing. **RCF:** conceptualization, funding acquisition, methodology, project administration, resources, supervision, writing – review and editing.

## Supplementary materials

**Supplementary Table 1: Full list of online-bioinformatics tools used during the workshop**

**Supplementary Table 2: Workshop agenda**

## Notes

### Competing Interest Statement

The authors have declared no competing interest.

## References

1. O’Neill J. Tackling Drug-resistant Infections Globally: Final Report and Recommendations. Review on Antimicrobial Resistance; 2016.

2. Murray CJL, Ikuta KS, Sharara F, Swetschinski L, Robles Aguilar G, Gray A, et al. Global burden of bacterial antimicrobial resistance in 2019: a systematic analysis. The Lancet. 2022;399(10325):629–55.

3. Hendriksen RS, Bortolaia V, Tate H, Tyson GH, Aarestrup FM, McDermott PF. Using Genomics to Track Global Antimicrobial Resistance. Frontiers in public health. 2019;7:242-.

4. Eyre DW. Infection prevention and control insights from a decade of pathogen whole-genome sequencing. J Hosp Infect. 2022;122:180–6.

5. Ishack S, Lipner SR. Bioinformatics and immunoinformatics to support COVID-19 vaccine development. J Med Virol. 2021;93(9):5209–11.

6. Sunita, Sajid A, Singh Y, Shukla P. Computational tools for modern vaccine development. Hum Vaccin Immunother. 2020;16(3):723–35.

7. Christoffels A, Mboowa G, van Heusden P, Makhubela S, Githinji G, Mwangi S, et al. A pan-African pathogen genomics data sharing platform to support disease outbreaks. Nat Med. 2023;29(5):1052–5.

8. World Health Organization (WHO). GLASS whole-genome sequencing for surveillance of antimicrobial resistance.. Geneva: World Health Organization; 2020.

9. Mlotshwa BC, Mwesigwa S, Mboowa G, Williams L, Retshabile G, Kekitiinwa A, et al. The collaborative African genomics network training program: a trainee perspective on training the next generation of African scientists. Genetics in Medicine. 2017;19(7):826–33.

10. Karikari TK. Bioinformatics in Africa: The Rise of Ghana? PLOS Computational Biology. 2015;11(9):e1004308.

11. Omotoso OE, Teibo JO, Atiba FA, Oladimeji T, Adebesin AO, Babalghith AO. Bridging the genomic data gap in Africa: implications for global disease burdens. Globalization and Health. 2022;18(1):103.

12. Shaffer JG, Mather FJ, Wele M, Li J, Tangara CO, Kassogue Y, et al. Expanding Research Capacity in Sub-Saharan Africa Through Informatics, Bioinformatics, and Data Science Training Programs in Mali. Front Genet. 2019;10:331.

13. Gurwitz KT, Aron S, Panji S, Maslamoney S, Fernandes PL, Judge DP, et al. Designing a course model for distance-based online bioinformatics training in Africa: The H3ABioNet experience. PLoS computational biology. 2017;13(10):e1005715–e.

14. Ras V, Botha G, Aron S, Lennard K, Allali I, Claassen-Weitz S, et al. Using a multiple-delivery-mode training approach to develop local capacity and infrastructure for advanced bioinformatics in Africa. PLoS computational biology. 2021;17(2):e1008640–e.

