## Supplementary Material for "Enable, Empower, Succeed: Harnessing Open Science for Antimicrobial Resistance Containment"

**Supplementary materials**

**Supplementary Table 1: Full list of online-bioinformatics tools used during the workshop**

| **Bioinformatics Step** | **Tools and Databases** | **Summary objectives** | **Learning outcomes** | **Exercise/Activity** | **Competencies/Outcomes** |
| --- | --- | --- | --- | --- | --- |
| Quality Control | FASTQC | Analyse the quality of genomic data with FASTQC | - Perform quality check with FASTQC - Interpret FASTQC and MultiQC report | Exercise/Activity 1: Case-based learning Fastqc and Multiqc | Interpretation of data quality results |
| Alignment | NCBI BLAST | Sequence alignment theory and applications | Interpret the information conveyed in NCBI BLAST search outputs and infer their significance | Exercise/Activity 1: Workgroup on sequence alignment | Interpretation of sequence alignment analysis |
|  |  | BLAST theory and practice | Examine the annotations of reported matches and their provenance | Exercise 2: Practical BLAST with selected nucleotide query sequences |  |
|  |  |  | Export BLAST results | Exercise 3: Practical BLAST with selected protein query sequences |  |
| In Silico prediction | Comprehensive Antibiotic Resistance Database (CARD) | Explore web-based tools as a method to access publicly available genomes | Browse publicly available genomes and associated metadata. | Exercise/Activity 1: Practical session on CARD, RGI, AMRFinderPlus, and PathogenWatch | **Online-based analysis, interpretation of data quality and in silico outputs** |
|  | ResFinder, VirulenceFinder, PlasmidFinder, of the Center of Genomic Epidemiology |  | Use online tools to analyse genome sequence data |  |  |
|  | AMRFinderPlus | Analyse genomic data | Interpret output of *in silico* prediction of antibiotic resistance | Exercise 2: Bacterial pathogenicity and mobilization (VirulenceFinder and PlasmidFinder from the Centre of Genomic and Epidemiology, PathogenWatch) |  |
|  | PathogenWatch |  | Interpret output of in silico prediction of mobile genetic elements and virulence |  |  |
| Lineage identification and phylogenetic | MLST of the Center of Genomic Epidemiology | Principles of phylogenetics | Understand the key principles of phylogeny | Exercise/Activity 1: Discussion on MicroReact phylogenetic tree | **Online-based analysis and interpretation of phylogenetic tree** |
|  | MicroReact |  | Use BV-BRC for retrieval of available genome |  |  |
|  | Bacterial and Viral Bioinformatics Resource Center (BV-BRC; formerly PATRIC) |  | Construct a tree with CSIPhylogeny | Exercise 2: Practical session on CSIPhylogeny |  |
|  | CSIPhylogeny |  | Visualise a phylogenetic tree with MicroReact |  |  |
|  | PathogenWatch |  |  |  |  |
|  | FigTree | Phylogenetic tree construction, interpretation and visualisation | Interpret a phylogenetic tree | Exercise 3: Practical session on Figtree and MicroReact |  |

**Supplementary Table 2: Workshop agenda**

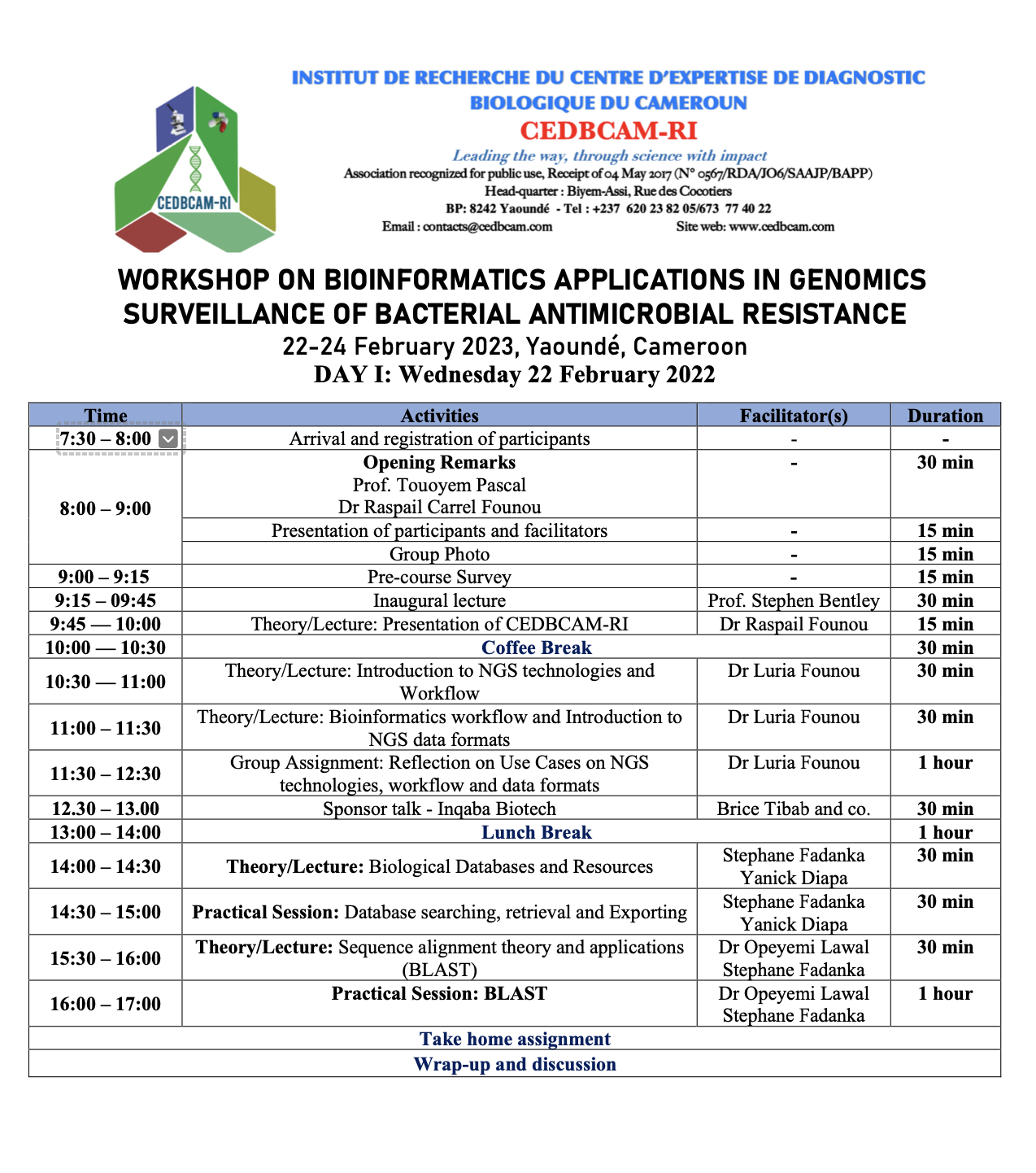

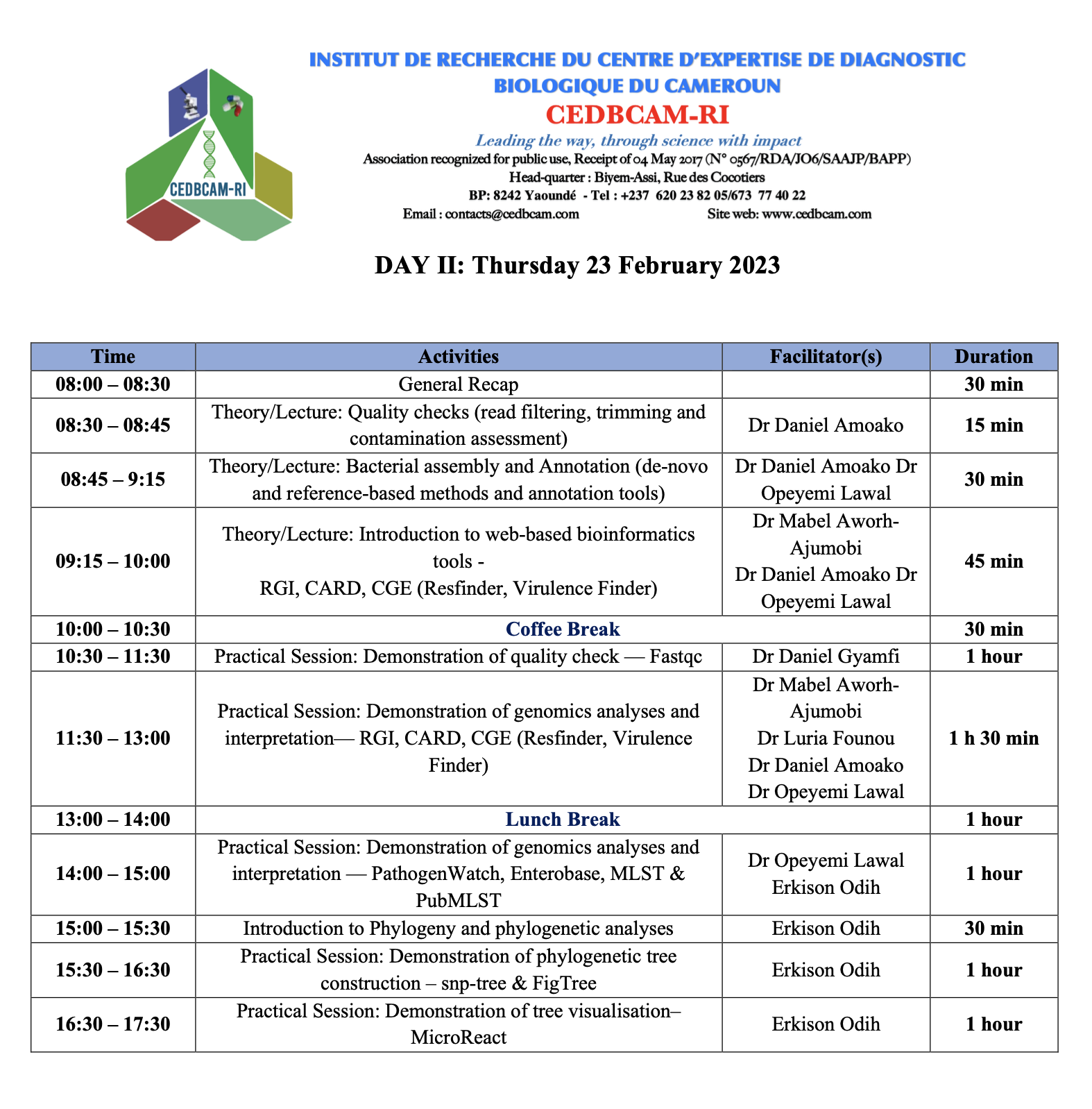

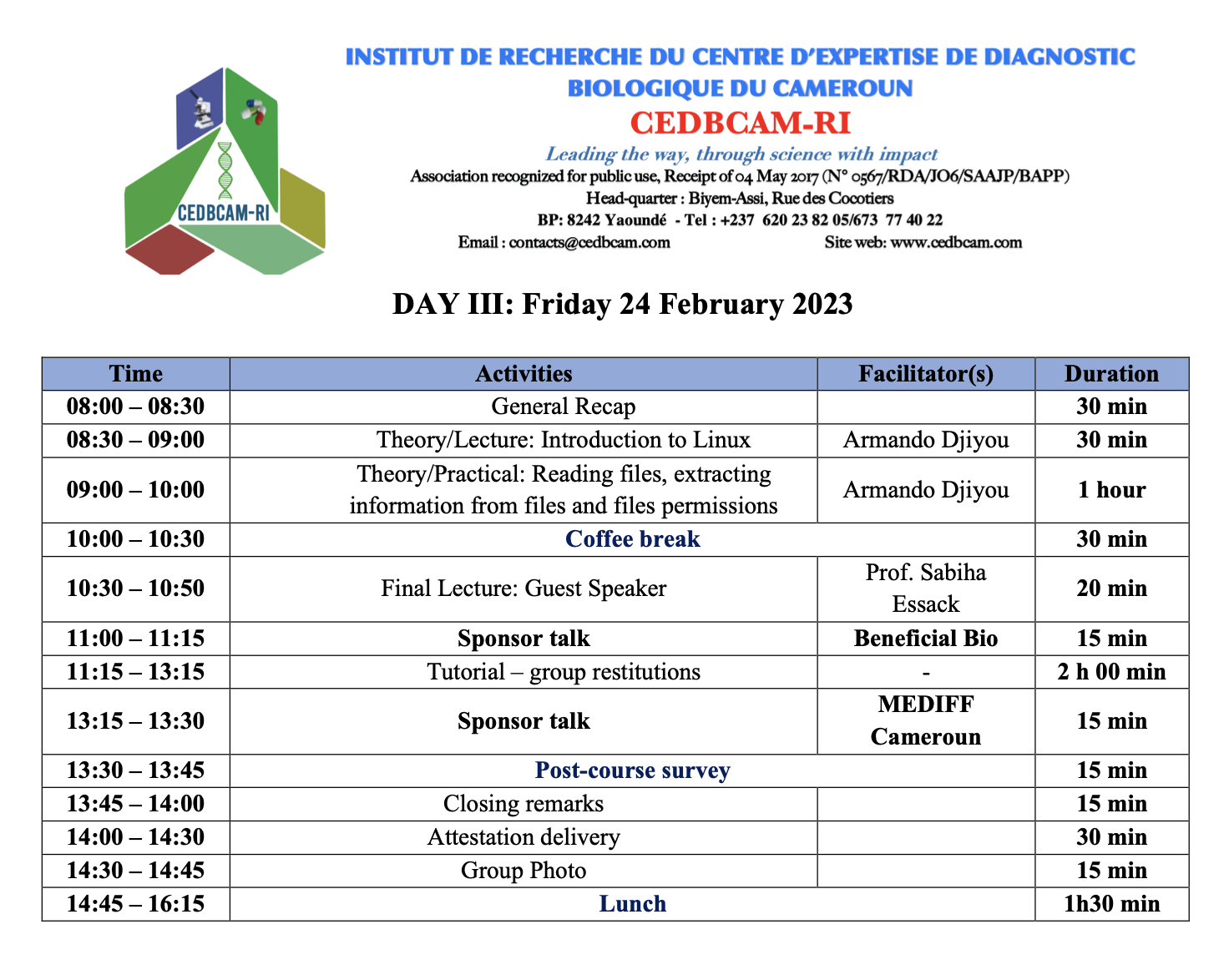
